# Transmural unipolar electrogram morphology is achieved within 7s at the posterior left atrial wall during pulmonary vein isolation: VISITAG™ Module-based lesion assessment during radiofrequency ablation

**DOI:** 10.1101/234799

**Authors:** David R. Tomlinson, Madison Myles, Kara N. Stevens, Adam J. Streeter

**Affiliations:** Dorset Heart Centre, Royal Bournemouth and Christchurch Hospitals NHS Foundation, Trust, Castle Lane East, Bournemouth, Dorset, BH7 7DW, UK.; Department of Medical Statistics, Plymouth University Peninsula Schools of Medicine and Dentistry, ITTC Building 1, Plymouth Science Park, Plymouth, Devon PL6 8BX, UK.

**Author notes:** South West Cardiothoracic Centre, Terence Lewis Building, Plymouth Hospitals NHS Trust, Plymouth, Devon, PL6 8DH, UK.

**Keywords:** Atrial fibrillation, contact force catheter ablation, pulmonary vein isolation, unipolar electrogram, VISITAG™ Module

## Abstract

**Aims:** To assess the occurrence of a histologically validated measure of transmural (TM) atrial ablation – pure R unipolar electrogram (UE) morphology change – at first-ablated left atrial posterior wall (LAPW) sites during contact force (CF)-guided pulmonary vein isolation (PVI).

**Methods and results:** Exported VISITAG™ Module and CARTOREPLAY™ (Biosense Webster Inc.) UE morphology data was retrospectively analysed in 23 consecutive patients undergoing PVI under general anaesthesia. PVI without spontaneous / dormant recovery was achieved in all, employing 16.3[3.2] minutes (mean [SD]) of temperature-controlled RF at 30W. All first-ablated LAPW sites demonstrated RS UE morphology pre-ablation, with RF-induced pure R UE morphology change in 98%. Time to pure R UE morphology was significantly shorter at left-sided LAPW sites (4.9[2.1] s versus 6.7[2.5] s; p=0.02), with significantly greater impedance drop (median 13.5Ω versus 9.9Ω; p=0.003). Importantly, neither the first-site RF duration (14.9 versus 15.0s) nor the maximum ablation catheter tip distance moved (during RF) were significantly different, yet the mean CF was significantly higher at right-sided sites (16.5g versus 11.2g; p=0.002). Concurrent impedance and objectively annotated bipolar electrogram (BE) data demonstrated ~6-8Ω impedance drop and ~30% BE decrease at the time of first pure R UE morphology change.

**Conclusion:** Using objective ablation site annotation, TM UE morphology change was typically achieved within 7s at the LAPW, with significantly greater ablative effect evident at left-sided sites. The methodology described in this report represents a novel and scientifically more rigorous foundation towards future research into the biological effects of RF ablation *in vivo*.

**Condensed abstract:** Through appropriate use of the VISITAG™ Module and CARTOREPLAY™, unipolar electrogram morphology change indicative of histologically confirmed transmural atrial ablation in animal models, was proven to occur typically within 7s, during first-site contact force-guided ablation at the left atrial posterior wall during pulmonary vein isolation.

## Introduction

Radiofrequency (RF) catheter ablation for atrial fibrillation (AF) is a complex undertaking, since each lesion must be transmural (TM) and irreversible, yet without incurring excessive risk of extra-cardiac thermal trauma. Considerable research efforts have focussed on the identification of optimal targets for lesion creation during pulmonary vein isolation (PVI) and following the development of contact-force (CF) sensing catheters, recommendations for the CF “working range” and force time integral (FTI) were derived.^1,2^ However, the recent report of a 6-fold greater proportion of atrio-oesophageal fistula (AOF) cases associated with CF-sensing catheter use compared to non-CF sensing catheters from a FDA MAUDE database analysis is concerning and suggests that these recommendations may be inappropriate.^3^

It should be acknowledged that this entire evidence-base for target RF delivery during PVI was derived using subjective means of ablation catheter stability assessment. In TOCCATA (the first study reporting a relationship between CF and clinical outcome following PVI), valid ablation records for analysis were those “with stable CF for at least 15 consecutive seconds”.^1^ However, for this methodology to be appropriate two assumptions must be satisfied, neither validated *in vivo:* (1) CF “stability” (however that is defined) must equate with a certain “fixed” degree of catheter tip-tissue positional stability; (2) TM ablation during PVI must never be achieved within 15s. Therefore, these data should be considered provisional, pending further investigations following appropriate methodological advances.

From a theoretical perspective, greater understanding of the RF energy required to achieve TM lesions during PVI requires novel methodology to define a stable point of RF delivery within the human heart through entirely objective annotation processes, and accurate means to assess the TM nature of the tissue response to RF at any such site. Logically, one fundamental element of ablation catheter positional stability is constant (versus intermittent) catheter-tissue contact. Adding complexity, both respiration and cardiac cycle-induced motion may create displacement at the catheter-tissue interface even in the setting of constant contact. While this may be accounted for using either respiratory or cardiac cycle RF annotation adjustment (i.e. no displacement assumed if the catheter returns to consecutive end-expiratory / end-diastolic sites within a pre-defined distance range), this may introduce error since RF delivery can only be proven to be “single site” when measures of catheter-tissue interaction and tissue response to RF at sites *without* catheter displacement are defined, and shown to be present at sites demonstrating respiration or cardiac cycle-induced motion yet “adjusted” to a single site. Therefore, at least until any such “motion adjustment” algorithms are proven to be accurate, avoidance of respiration / cardiac cycle adjustment should provide the most appropriate methodology underlying objective annotation of stable ablation catheter positions *in vivo*.

When considering the tissue response to RF application, electrogram changes indicating TM atrial ablation effects are well described in animal studies. Otomo *et al* identified that “pure R” unipolar electrogram (UE) morphology was 100% predictive of histologically proven TM lesions in porcine atria, and was catheter-tissue angle independent.^4^ Using a contact-force (CF) sensing catheter in dogs, Bortone *et al* demonstrated that 95% of atrial lesions were histologically TM following immediate RF termination at the first occurrence of a pure R UE, with no evidence of extra-cardiac thermal trauma. However, continuous RF delivery for 10 or 20s beyond this time (and indeed a “conventional” 30s duration application) consistently resulted in 100% TM lesions yet with 11-17% lesions associated with extracardiac thermal trauma.^5^ Considering the employed power-controlled RF protocol at 30W (48°C, 17ml/min irrigation) and minimum CF 10g resulted in pure R UE morphology at 7s and with mean lesion depth of 4.3mm, one would anticipate a high risk of extra-cardiac thermal trauma from the practice of 20-40s / “a median of 45s” RF “per site” during recently conducted CF-guided PVI studies in humans.^6,7^

Here we provide the first-in-man report of UE morphology changes at sites of first RF application on the left atrial posterior wall (LAPW) during first-time PVI, employing an objective method for ablation catheter stability annotation fulfilling the criteria for a stable point of RF delivery *in vivo* outlined above, via automated annotation software (VISITAG™ Module with CARTOREPLAY™, Biosense Webster Inc., Diamond Bar, CA). In addition, we report concurrent and objectively annotated data regarding bipolar electrogram (BE) attenuation and impedance drop. Finally, through extended catheter tip positional data analysis using the software R^8^, we report the relationship between measured catheter tip displacement during RF, time to pure R UE morphology and impedance drop at annotated LAPW ablation sites. This ablation protocol and the technical approach to PVI was developed during an Institutional Review Board (IRB)-approved service evaluation, and was without influence from any published recommendations for RF delivery derived from studies employing subjective methods of ablation catheter stability assessment; a complete description can be found online.^9^

## Methods

Retrospective analysis of VISITAG™ Module and CARTOREPLAY™ data was performed following single-operator PVI, in consecutive unselected patients with symptomatic AF undergoing PVI according to current treatment indications.^10^ Briefly, all procedures were undertaken using general anaesthesia (GA) with endotracheal intubation and intermittent positive pressure ventilation. Single transseptal access was obtained with an SL1 (St Jude Medical Inc., Minneapolis, MN) sheath, following which either a NaviStar^®^ THERMOCOOL^®^ SMARTTOUCH™ (ST) F curve or NaviStar^®^ EZSTEER^®^ THERMOCOOL^®^ SMARTTOUCH™ D/F catheter (Biosense Webster) via an Agilis™ NxT sheath (St Jude Medical Inc.) was placed in the left atrium (LA) via the first transseptal site. ACCURESP™ Module (Biosense Webster) respiratory training was checked with a duodecapolar LASSO^®^ Nav catheter (Biosense Webster) in both the right superior pulmonary vein (PV) and the left inferior PV and applied prospectively as required to the CARTO^®^3 geometry (V.3, Biosense Webster).

CF-guided PVI was performed in temperature-controlled mode (17ml/min, 48°C) at 30W, with lesion placement guided by the VISITAG™ Module.^9^ The preferred sites of first RF application were at the LAPW opposite each superior PV ~1cm from the PV ostium, although in cases where constant catheter-tissue contact could only be achieved with maximal CF ≥70g an adjacent LAPW site with lower peak CF was chosen; the target first-site ablation time was 15s. Circumferential PVI (entrance and exit block) was thereafter achieved with continuous RF delivery, with additional point-by-point RF applied as required to achieve target parameters. Spontaneous recovery of PV conduction was assessed and eliminated during a minimum 20-minute wait. Dormant recovery was evaluated and targeted a minimum of 20 minutes after the last RF.

### VISITAG™ Module and CARTOREPLAY™ use

The VISITAG™ Module permits automated ablation site annotation through catheter “stability filters” utilising CF and positional data during RF application. The ablation catheter tip position is measured every 16/17ms, from which the standard deviation (SD) is calculated over a minimum interval of 60 positions (i.e. 1s). Consecutive ablation catheter tip positions within a user-defined maximum range for the SD (in mm) during on-going RF are used to annotate a site meeting positional stability filter requirements. CF is measured every 50ms, with the CF filter employing a user-defined minimum CF for a percentage of RF time at sites meeting positional stability filter requirements. Consecutive catheter tip positions satisfying both positional and CF stability filter requirements beyond the minimum user-defined duration (system minimum, 3s) result in automated ablation annotation; a 3D and/or surface-projected tag may be used to denote the mean catheter position at each site. Post-ablation, annotated tags provide summative ablation data including RF duration, mean and range of CF, impedance and FTI (figure 1A and B; see box, left panels), while an export function permits extended data analysis. Following completion of a service evaluation (July – December 2013),^9^ the following VISITAG™ Module filter settings were used during this present report: Positional stability duration 3s and range 2mm; force-over-time 100% minimum 1g (i.e. constant catheter-tissue contact). Regardless of the findings from ACCURESP™ Module respiration-induced motion assessment, respiration adjustment was never applied to the VISITAG™ Module filter settings. For extended ablation catheter positional stability analysis, R software code was written to analyse exported VISITAG™ Module data regarding catheter tip (position) SD and maximum displacement from the mean position annotated at each first site of LAPW ablation. Impedance and RF power data at 100ms intervals during RF were obtained via the VISITAG™ Module export function.

**Figure 1:**
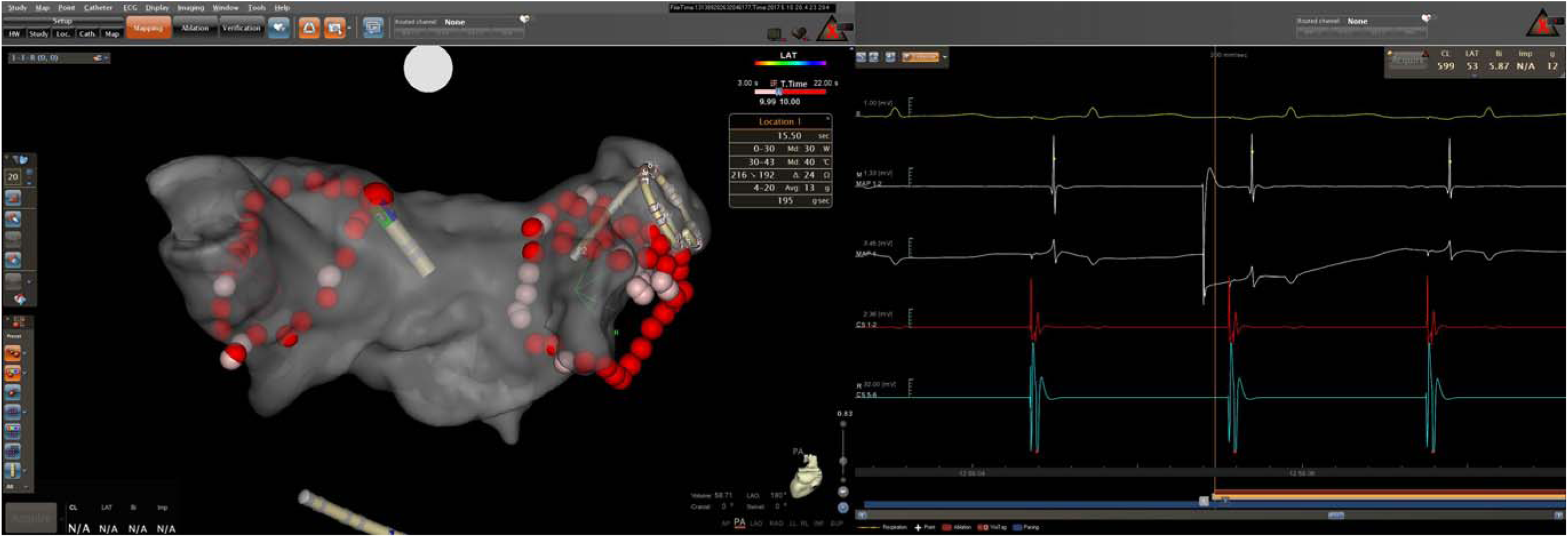
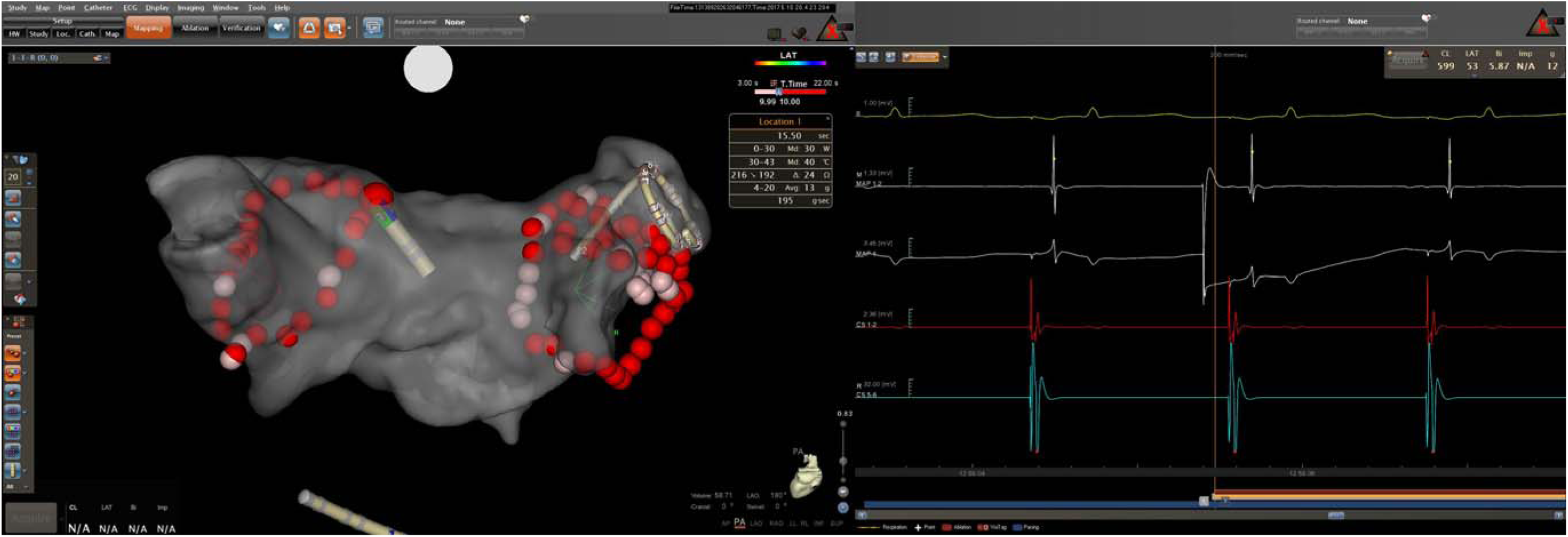
Left panel showing CARTO^®^3 LA geometry with VISITAG™ Module ablation site annotation displayed as 2mm radius spheres, with transition to red at 10s RF duration (transparent PA view) and right panel showing CARTOREPLAY™ screen with 24-hour clock timeline beneath. A: First RF application annotation onset at the LAPW during left PV isolation. “Location 1” is highlighted (greater tag diameter) with corresponding annotated summative ablation site data shown (box). CARTOREPLAY™ electrogram data at 200ms speed demonstrates pre-ablation RS UE morphology during CS pacing at 600ms cycle length (from top to bottom: ECG lead II; MAP 1-2 bipolar electrogram 1.33mV scale; MAP 1 UE 3.45mV scale; CS 1-2 and 5-6 bipolar electrograms). RF and VISITAG™ Module annotation onset is identified in the middle of the CARTOREPLAY™ screen as white and red/yellow horizontal bars at the bottom, respectively. B: CARTO^®^3 LA geometry and CARTOREPLAY™ screen showing time calliper at the onset of 3 consecutive pure R UE morphology complexes during CS pacing at “Location 1”, following 3.1s RF.

CARTOREPLAY™ provides a means to retrospectively analyse electrogram data at all VISITAG™ Module annotated ablation sites, within 18 hours (data is deleted thereafter). During this present report, UE morphological analysis was performed immediately following case completion, employing histologically validated reported criteria for TM RF effect (i.e. pure R morphology change). UE filtering was 0.5-120Hz with analysis display speed 200mm/s and voltage scale 3.45 or 3.80mV. The time to pure R UE morphology was taken at the first of 3 consecutively occurring pure R complexes, no more than 1 of which could be an atrial ectopic. In order to facilitate summative analysis for all first-ablated LAPW sites, every RF application was performed during CS pacing at 600ms cycle length. Ablation catheter distal bipolar electrogram (BE) peak-to-peak amplitude was obtained using automated Wavefront Annotation (Biosense Webster). Pictorial evidence of the stated UE morphological changes was provided via a JPEG creation tool (e.g. figures 1A and B); all first-ablated LAPW site images are shown in the data supplement.

Statistical analyses were performed using GraphPad Prism version 4.03. Parametric data are expressed as mean [SD]; non-parametric data are presented as median (1st – 3rd quartile). Unpaired / paired t test or the Mann Whitney test was used to assess statistical significance for continuous data, as appropriate. P < 0.05 indicated a statistically significant difference. This work received IRB approval for publication as a retrospective service evaluation; all patients provided written, informed consent.

## Results

Twenty-three patients underwent first-time PVI as described, between 1^st^ December 2016 and 11^th^ May 2017: 12 patients had persistent AF, 11 PAF; 17 were male (74%); mean age 58 [13] years and CHA_2_DS_2_-VASc score 1.4 [1.4]. Complete PVI was achieved in all without spontaneous / dormant recovery of PV conduction, following 16.3 [3.2] minutes of RF, with no procedural complications.

Pre-ablation, all first-ablated LAPW annotated sites demonstrated RS UE morphology (e.g. figure 1A and data supplement). For sites adjacent to the left PV, ablation-induced pure R UE morphology was identifiably achieved in 21/23 cases (e.g. figure 1B and data supplement). One case of inadvertent catheter displacement resulted in annotation termination at 4.43s when RS UE morphology remained present (figure 5A, data supplement), while in another, gross artefact made UE morphology assessment impossible (figure 15A, data supplement). For annotated sites adjacent to the right PV, ablation-induced pure R UE morphology was achieved in all. However, one case of inadvertent catheter displacement at RF onset resulted in 4.97s non-annotated RF delivery, so the time to pure R UE measurement was deemed invalid for analysis.

Annotated first-ablated LAPW site data is shown in table 1. At left-sided sites there was a significantly shorter time to pure R UE morphology (4.9[2.1] s versus 6.7[2.5] s; p=0.02), with a significantly greater total impedance drop (13.5Ω versus 9.9Ω; p=0.003). This was not explained on the basis of a difference in the total RF duration, while both the mean CF and FTI were significantly lower for left-sided sites (11.2g versus 16.5g; p=0.002 and 167gs versus 244gs; p=0.004). Furthermore, while the maximum ablation catheter tip distance moved from the (annotated) mean point was not significantly different, the standard deviation of catheter tip motion was significantly greater at left-sided sites (1.4[0.3] mm versus 1.1[0.3] mm; p=0.001). There were no significant differences between the BE (distal ablation catheter pair) data, although there was a trend towards greater total percentage BE decrease at leftsided sites (61[19] % versus 51[24] %).

**Table 1:**
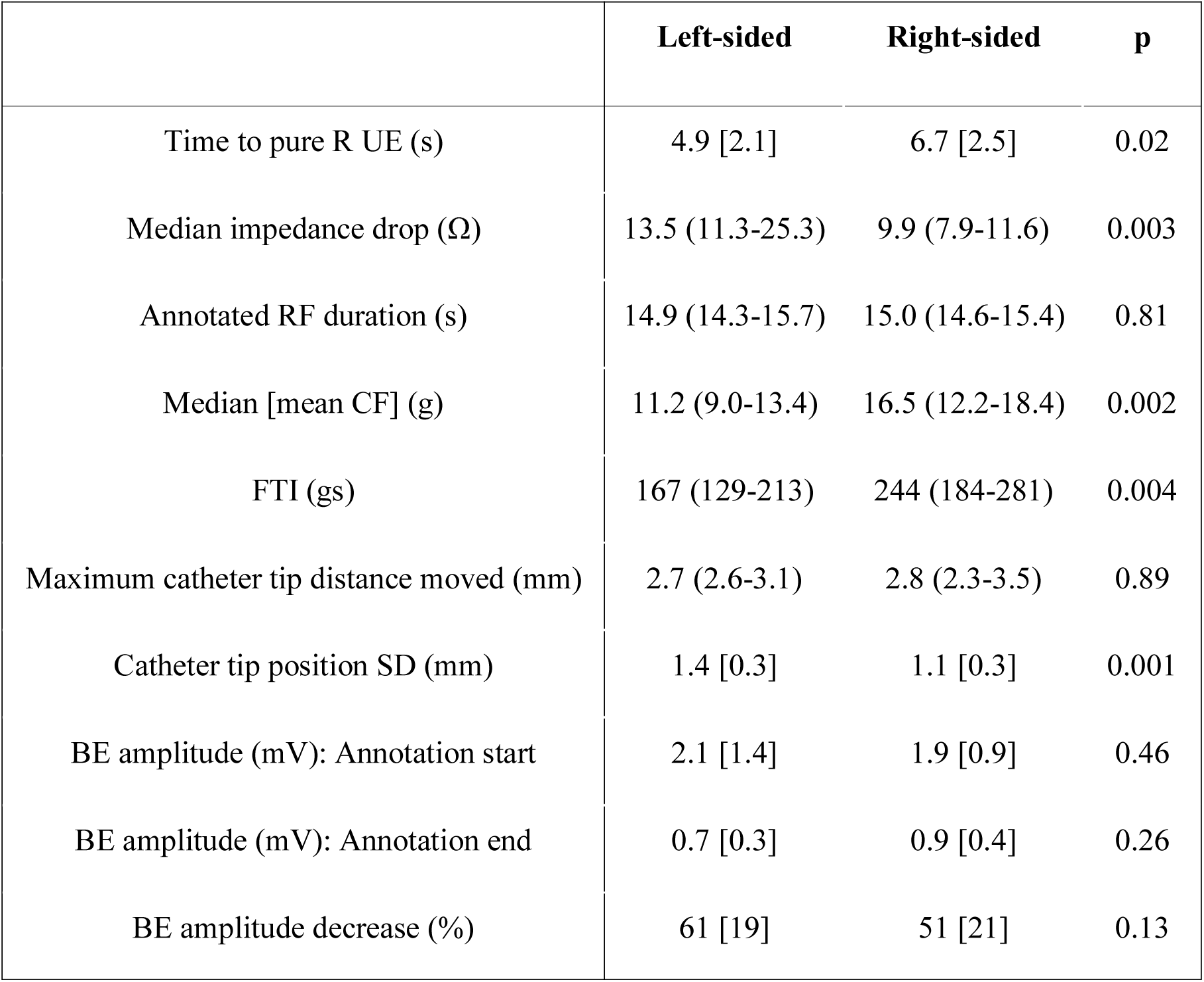
Left atrial posterior wall first-ablated site automated annotation data, according to proximity to the left or right pulmonary vein (left-sided / right-sided, respectively). For bipolar electrogram (BE) data, annotation “start” and “end” represent the first and last ablation catheter (distal pair) electrograms respectively, during automated VISITAG™ Module RF annotation. Data is shown as mean [SD] / median (25^th^ – 75^th^ centile), as appropriate. UE, unipolar electrogram; CF, contact force; FTI, force time integral; SD, standard deviation.

The first 15s of annotated impedance and percentage bipolar electrogram decrease data is shown in figure 2, with RF power below. At the times corresponding to the pure R UE morphology data in table 1, an impedance decrease of ~6-8Ω can be observed, accompanied by ~30% BE decrease for both left-sided and right-sided ablation sites. Notably, utilising this temperature-controlled RF protocol the delay to achieve 30W was ~6s. Correlation between the time to pure R UE morphology and the following factors was assessed: Total impedance drop; mean CF; maximum catheter tip distance moved; ablation catheter tip position SD; and BE amplitude (annotation start). The only significant correlation was in right-sided sites and with impedance drop – Spearman r −0.43 (−0.72 to −0.005; p=0.048).

**Figure 2:**
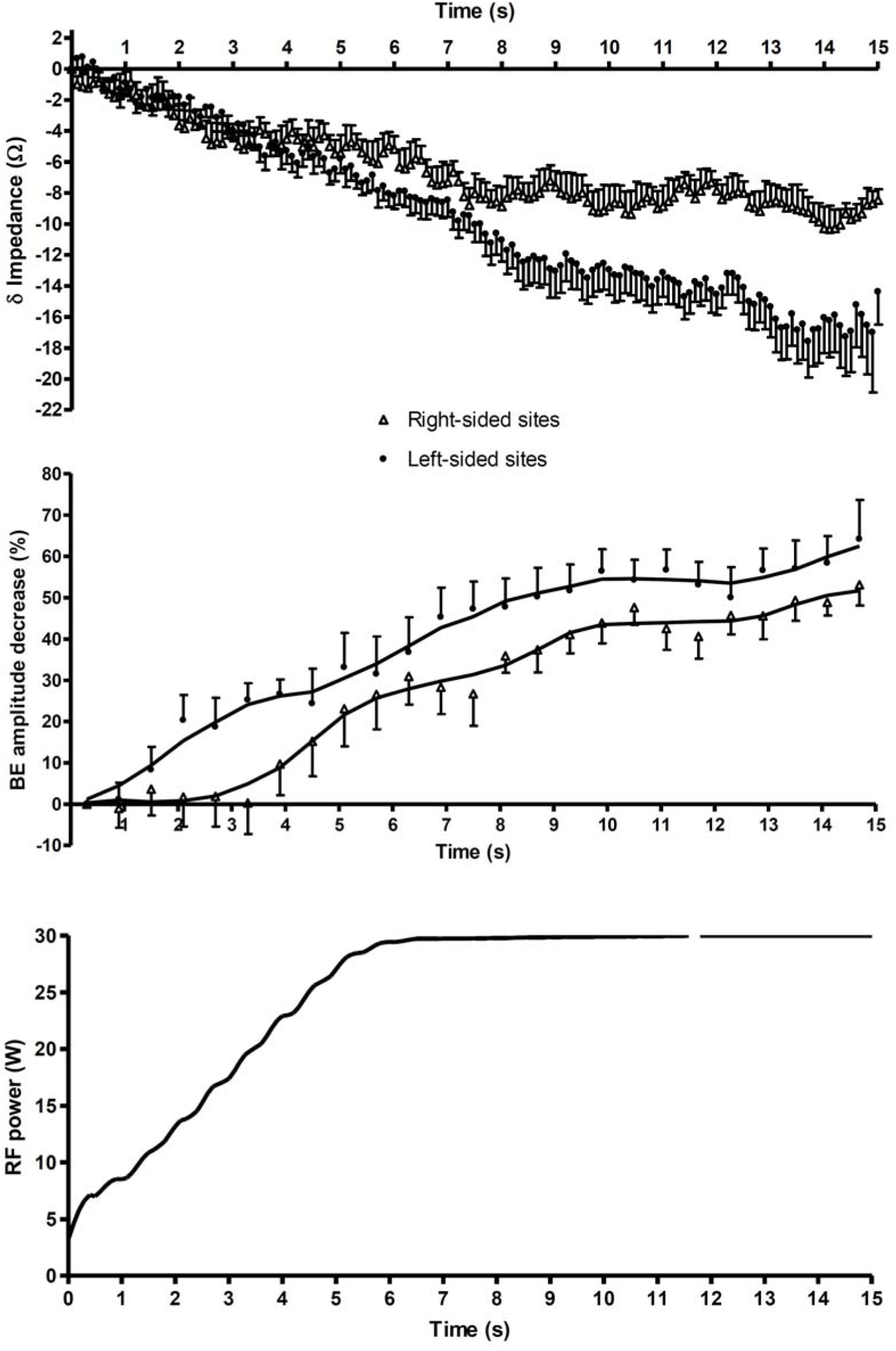
Impedance change and percentage bipolar electrogram decrease (ablation catheter distal pair) for the first 15s of annotated ablation site data according to proximity to the left or right PV (left-sided / right-sided, respectively). Error bars are shown in one direction for clarity. The RF power data represents a smoothed curve from all RF applications.

## Conclusions

A key problem for any operator performing CF-guided PVI is successfully striking a balance between the competing risks of arrhythmia recurrence due to incomplete lesion formation^11^ and extra-cardiac thermal trauma due to excessive local RF energy delivery.^3^ Indeed, due to the close proximity of the oesophagus to the LAPW, inadvertent oesophageal thermal injury may be seen in up to 48% patients^12^. Worryingly, reports of an increase in the rate of atrio-oesophageal fistula following the introduction of CF-sensing catheters^3,13^ indicates a likely causative relationship and highlights the risk of conducting PVI with an imperfect understanding of when TM ablation has been achieved.

The analyses described in this present report represent the first-in-man description of the LAPW “physiological response” to RF application at objectively annotated first-ablated sites meeting a suitable definition of a stable point of catheter tip-tissue contact in 3D space. Specifically, we have shown that when an ablation catheter is maintained within a (SD) range of 2mm, acutely durable ablation effect was universally achieved following 15s temperature-controlled RF at 30W, with TM UE morphology change typically at <7s associated with a BE attenuation of ~30% and impedance drop of ~6-8Ω. Furthermore, we have identified superior catheter tip-tissue energy coupling for left-sided ablation sites, evidenced by significantly shorter time to pure R UE morphology, associated with a significantly greater total impedance drop. Finally, the absence of a significant correlation between the time to pure R UE morphology change and either the maximum catheter tip distance moved / tip position SD provides supportive evidence for the appropriateness of the 2mm SD positional stability filter “working range”, without ACCURESP™ adjustment annotation protocol.

That these data are out of keeping with current views regarding what constitutes a “conventional / suitable” approach to RF application during PVI should come as no surprise when considering the development of the current approach towards PVI and evidence available from other publications: (1) The original description of ~30s RF “per site”, aiming for 80% BE attenuation was without scientific validation,^14^ yet has remained unchallenged and perpetuated in methodology ever since;^6,15^ (2) the methods used to derive current CF and FTI recommendations are flawed,^1,2^ compared to investigations conducted utilising objective means of ablation catheter stability assessment; (3) the LA wall is thin (e.g. computed tomography assessment pre-ablation mean thickness 1.9mm; range 0.5-3.5mm),^16^ yet laboratory preparations of irrigated RF using “conventional” RF duration result in far deeper lesions (e.g. in a bovine skeletal muscle model of intermittent catheter contact at 20W, an FTI of 200gs (CF range 0-22g) resulted in 4mm depth lesions);^17^ (4) similarly and as previously stated, *in vivo* investigations of CF-guided ablation in canines demonstrated pure R UE morphology change at 7s, with histologically confirmed TM ablation effect in 95% applications (power-controlled RF at 30W), and mean lesion depth 4.3mm;^5^ (5) a report of real-time lesion assessment utilising a novel combined ultrasound and RF ablation catheter *in vivo* in sheep, demonstrated rapid lesion development during right atrial epicardial ablation at 7W (i.e. 1.5mm depth at 10s, see figure 2A, Wright *et al*2011);^18^ (6) oesophageal temperature alerts during LAPW ablation at 25W (target CF 10-20g), occurred in 73% patients, with intra-luminal temperature 39°C illustrated as early as 7s following RF onset (see figure 2, Halbfass *et al* 2017).^19^

The findings in this present report have numerous implications. Most importantly, we have demonstrated that the VISITAG™ Module and CARTOREPLAY™ may together be used as investigational tools during PVI, permitting an assessment of the TM nature of the tissue response to RF delivery. Clearly, this provides a foundation for further investigations in an effort to reduce the risk of oesophageal thermal injury (e.g. evaluating the acute effectiveness of 20W power to the LAPW). However, in view of the objective nature of VISITAG™ Module annotation, any operator may similarly investigate lesion creation utilising their “usual approach” to ablation during CF-guided PVI. Furthermore, in view of the described evidence of rapid TM ablation effect in thin-walled atrial tissue, such investigations are likely to improve the safety profile of CF-guided PVI, through a movement away from the conventional view of a requirement for ~30s RF delivery “per site”, achieving 80% BE^6,14,15^ attenuation and FTI 400gs,^1,2^ towards shorter RF duration and possibly lower power delivery. This represents an important advance in RF ablation research methodology, since when the VISITAG™ Module is used as described in this present report, it comes as close as is presently possible to representing a “universal language” for the RF ablation “effector arm”. Consequently, and through the consistent application of CF and positional stability filters described here, future research protocols may be optimised, practice harmonised and with published outcomes then replicated by any operator able to maintain the catheter position and CF stability in the manner described.

The methodology presented here also represents a departure from studies utilising automated ablation site annotation yet failing to recognise the limitations of a CF filter permitting intermittent catheter-tissue contact (i.e. 30% minimum 5g) and the non-validated use of ACCURESP™ adjustment,^6^ when considering a suitable definition for a stable point of RF application *in vivo*. Furthermore, an inherent additional problem with any system of ablation lesion delivery founded upon assessments of CF and energy delivery alone is clearly apparent when considering the differential ablative effect between left and right-sided sites demonstrated in this report. However, this should also not be surprising when recognising the recently reported range of the maximal impedance drop resulting from an ablation index (AI) value of 300; i.e. 0 to ~50Ω (see figure 3, Das *et al*)^6^

### Study limitations

All UE morphology data were collected retrospectively, so this report does not prove that modification of RF delivery based on real-time UE morphological assessment during VISITAG™ Module and CF-guided PVI is appropriate. However, this is worthy of further research, not least in view of the promising findings from a single-centre UE signal modification-based PVI study in humans, before the advent of CF-sensing and objective ablation site annotation.^20^ Furthermore, neither oesophageal luminal temperature monitoring nor post-ablation endoscopic evaluation was employed; given the time to pure R UE morphology change of 3.1s demonstrated in figure 1 it is highly likely that some LAPW lesions were accompanied by inadvertent extra-cardiac thermal injury. Therefore, the approach to ablation described in this report is not a direct recommendation for others to follow without careful consideration. Rather, through adopting the methodology described in this present report there is now the opportunity for all to identify the most appropriate means to achieve TM ablation during PVI without excessive risks, using insights into one’s “usual approach”, through objective ablation site annotation.

The significantly greater time to pure R UE morphology at right-sided sites may in part have been due to increased LA wall thickness. Also, although the maximum distance of catheter tip motion did not differ between left and right-sided sites, there may have been a greater degree of LAPW motion beneath the catheter tip at right-sided sites, despite a significantly greater mean CF (i.e. out-of-phase, lateral motion). However, the significantly greater total impedance drop at left-sided sites remains the first demonstration of superior regional catheter tip-tissue energy coupling *in vivo*. Although this finding cannot be fully explained at present, these data may indicate that the more oblique catheter orientation routinely obtained during left-sided ablation may result in a greater surface area of ablation electrode in contact with the tissue, with even a relatively small difference potentially resulting in significantly greater power delivery.^21^ Although the mechanisms underlying these observations await identification, this represents an important unknown variable in any attempt to harmonise RF ablation practice based upon ablation lesion quality markers incorporating CF, RF duration and power alone.^6^

Maintaining a stable catheter position within the chosen 2mm SD range may require some novel operator “adjustments” for respiration-induced motion, particularly when the use of ACCURESP™ adjustment use during PVI remains non-validated. There is presently no means to describe any compensatory sheath / catheter movements utilised; jet ventilation may therefore improve the reproducibility of VISITAG™ Module and CF-guided PVI, but this requires further study. Finally, achieving durable PVI requires sufficient overlap between adjacent lesions; these present data provide no information concerning the diameter of TM ablative effects. However, through conducting similar investigations to those described here, the diameter of RF-induced thermal injury may be identified during PVI; these data have been collected and will be the subject of a future report.

### Conclusions

Through appropriate use of the VISITAG™ Module and CARTOREPLAY™ during PVI, the TM nature of the tissue responses to RF may be investigated. These annotation tools provide a research foundation with greater scientific rigour, potentially better informing future RF ablation practice.

## Acknowledgments

I am grateful to Cherith Wood, Daniel Newcomb and Ian Lines, Cardiac Physiologists, for their technical support into all cases conducted in this report. I am also grateful to Robert Pearce and Vicky Healey (Biosense Webster Inc.) for additional technical assistance and to Noam Seker-Gafni, Tal Bar-on, Einav Geffen, Assaf Rubissa and colleagues at the Haifa Technology Center, Israel for their help with VISITAG™ Module technical queries.

## References

1. Reddy VY, Shah D, Kautzner J, Schmidt B, Saoudi N, Herrera C, et al. The relationship between contact force and clinical outcome during radiofrequency catheter ablation of atrial fibrillation in the TOCCATA study. Heart Rhythm. 2012;9(11):1789–95.

2. Neuzil P, Reddy VY, Kautzner J, Petru J, Wichterle D, Shah D, et al. Electrical reconnection after pulmonary vein isolation is contingent on contact force during initial treatment: results from the EFFICAS I study. Circ Arrhythm Electrophysiol. 2015;6(2):327–33.

3. Black-Maier E, Pokorney SD, Barnett AS, Zeitler EP, Sun AY, Jackson KP, et al. Risk of atrioesophageal fistula formation with contact force-sensing catheters. Heart Rhythm. 2017;14(9):1328–33.

4. Otomo K, Uno K, Fujiwara H, Isobe M IY. Local unipolar and bipolar electrogram criteria for evaluating the transmurality of atrial ablation lesins at different catheter orientations relative to the endicardial surface. Heart Rhythm. 2010;7(9):1291–300.

5. Bortone A, Brault-Noble G, Appetiti A, Marijon E. Elimination of the negative component of the unipolar atrial electrogram as an in vivo marker of transmural lesion creation: acute study in canines. Circ Arrhythm Electrophysiol. 2015;8(4):905–11.

6. Das M, Loveday JJ, Wynn GJ, Gomes S, Saeed Y, Bonnett LJ, et al. Ablation index, a novel marker of ablation lesion quality: prediction of pulmonary vein reconnection at repeat electrophysiology study and regional differences in target values. Europace. 2017;19(5):775–83.

7. Park C-I, Lehrmann H, Keyl C, Weber R, SChiebeling J, Allgeier J, et al. Mechanisms of Pulmonary Vein Reconnection After Radiofrequency Ablation of Atrial Fibrillation: The Deterministic Role of Contact Force and Interlesion Distance. J Cardiovasc Electrophysiol. 2015;25(7):701–8.

8. R: A language and environment for statistical computing. R Core Team, Foundation for Statistical Computing, Vienna, Austria. 2017; https://www.r-project.org/.

9. Tomlinson DR. Derivation and validation of a VISITAG^TM^-guided contact force ablation protocol for pulmonary vein isolation. bioRxiv [Internet]. 2017 Dec 13 [cited 2017 Dec 14];232694. Available from: https://www.biorxiv.org/content/early/2017/12/13/232694

10. Calkins H, Kuck KH, Cappato R, Brugada J, Camm AJ, Chen S-A, et al. 2012 HRS/EHRA/ECAS expert consensus statement on catheter and surgical ablation of atrial fibrillation: recommendations for patient selection, procedural techniques, patient management and follow-up, definitions, endpoints, and research trial design. Heart Rhythm. 2012;9(4):632–696.e21.

11. Ouyang F, Antz M, Ernst S, Hachiya H, Mavrakis H, Deger FT, et al. Recovered pulmonary vein conduction as a dominant factor for recurrent atrial tachyarrhythmias after complete circular isolation of the pulmonary veins: lessons from double Lasso technique. Circulation [Internet]. 2005 Jan 18 [cited 2015 Nov 12];111(2): 127–35.

12. Di Biase L, Saenz LC, Burkhardt DJ, Vacca M, Elayi CS, Barrett CD, et al. Esophageal capsule endoscopy after radiofrequency catheter ablation for atrial fibrillation: documented higher risk of luminal esophageal damage with general anesthesia as compared with conscious sedation. Circ Arrhythm Electrophysiol [Internet]. 2009 Apr 1 [cited 2017 Nov 20];2(2):108–12.

13. Medeiros De Vasconcelos JT, Filho S dos SG, Atié J, Maciel W, De Souza OF, Saad EB, et al. Atrial-oesophageal fistula following percutaneous radiofrequency catheter ablation of atrial fibrillation: the risk still persists. Europace. 2017;19(2):250–58.

14. Pappone C, Rosanio S, Oreto G, Tocchi M, Gugliotta F, Vicedomini G, et al. Circumferential radiofrequency ablation of pulmonary vein ostia: A new anatomic approach for curing atrial fibrillation. Circulation [Internet]. 2000 Nov 21 [cited 2017 Nov 20];102(21):2619–28.

15. Arentz T, Weber R, Burkle G, Herrera C, Blum T, Stockinger J, et al. Small or Large Isolation Areas Around the Pulmonary Veins for the Treatment of Atrial Fibrillation?: Results From a Prospective Randomized Study. Circulation [Internet]. 2007 Jun 19 [cited 2017 Nov 20];115(24):3057–63.

16. Beinart R, Abbara S, Blum A, Ferencik M, Heist K, Ruskin J, et al. Left atrial wall thickness variability measured by CT scans in patients undergoing pulmonary vein isolation. J Cardiovasc Electrophysiol. 2011;22(11):1232–6.

17. Shah DC, Lambert H, Nakagawa H, Langenkamp A, Aeby N, Leo G. Area under the real-time contact force curve (force-time integral) predicts radiofrequency lesion size in an in vitro contractile model. J Cardiovasc Electrophysiol [Internet]. 2010 Sep [cited 2015 Nov 11];21(9): 1038–43.

18. Wright M, Harks E, Deladi S, Suijver F, Barley M, van Dusschoten A, et al. Real-time lesion assessment using a novel combined ultrasound and radiofrequency ablation catheter. Heart Rhythm [Internet]. 2011 Feb [cited 2017 Nov 15];8(2):304–12.

19. Halbfass P, Müller P, Nentwich K, Krug J, Roos M, Hamm K, et al. Incidence of asymptomatic oesophageal lesions after atrial fibrillation ablation using an oesophageal temperature probe with insulated thermocouples: a comparative controlled study. Europace. 2017;19(3):385–91.

20. Bortone A, Appetiti A, Bouzeman A, Maupas E, Ciobotaru V, Boulenc J-M, et al. Unipolar signal modification as a guide for lesion creation during radiofrequency application in the left atrium: prospective study in humans in the setting of paroxysmal atrial fibrillation catheter ablation. Circ Arrhythm Electrophysiol [Internet]. 2013 Dec [cited 2015 Nov 11];6(6):1095–102.

21. Wittkampf FHM, Nakagawa H. RF Catheter Ablation: Lessons on Lesions. Pacing Clin Electrophysiol [Internet]. 2006 Nov [cited 2017 Nov 20];29(11):1285–97.

